# Influence of advanced footwear technology spikes on middle- and long-distance running performance measures in trained runners

**DOI:** 10.1101/2024.04.13.589345

**Authors:** Víctor Rodrigo-Carranza, Violeta Muñoz de la Cruz, Wouter Hoogkamer

## Abstract

Two types of track spikes are commonly used, recently: spikes with a compliant and resilient midsole foam (e.g., PEBA), and spikes that combine such modern foam with a carbon fiber plate. Here we evaluated the effect of these different spike technologies on running performance measures for middle- and long-distance track events in trained runners. Fourteen females performed a single visit with six 200m trials at self-perceived 800m race pace in three different spike conditions (Control, PEBA and PEBA+Plate) twice in a mirrored order. Sixteen males completed four visits. During the first three visits they performed six 200m trials at self-perceived 800m race pace, twice in each condition. Subsequently, they performed a 3,000m time trial in one of the three spike conditions. During visit four, participants completed six 4-minute running economy trials at 5 m/s, twice in each condition. At 800m race pace females ran faster in PEBA (2.1%) and PEBA+Plate (2.0%) compared to Control. Males ran faster in PEBA (1.4%) and PEBA+Plate (2.4%) compared to Control, and in PEBA+Plate compared to PEBA (1.1%). Similarly, males ran the 3,000m time trial faster in PEBA (1.0%) and PEBA+Plate (2.4%) than in Control. Running economy was better in PEBA (5.1%) and PEBA+Plate (4.0%) than in Control. Performance benefits from modern foam spikes with and without a plate are equivalent for female middle-distance running, and for male long-distance running, but larger from modern foam spikes with a plate for male middle-distance running.

## INTRODUCTION

Since the winter of 2019, advanced footwear technology (AFT) spikes have significantly impacted performances in middle and long-distance track events, from 800m to 10,000m, both indoors and outdoors, for both males and females (Healey et al., 2022; Willwacher et al., 2024). Two different types of AFT spikes are commonly used: spikes with a modern compliant and resilient midsole foam [e.g., polyether block amide (PEBA)], and spikes that combine such modern foam with an element to increase longitudinal bending stiffness, often a carbon fiber plate. Most medals in the individual middle-distance (800m, 1,500m, 3,000m steeple chase) and long-distance track events (5,000m and 10,000m) for males and females at the 2021 Tokyo Olympics Games, were won by athletes wearing AFT spikes, with each type of AFT spike equally often used by medalist across these events.

It has been shown that AFT road shoes improve running economy (RE) (Hoogkamer et al., 2018; Barnes & Kilding, 2019) and long-distance running performance in events from the 10km to the marathon (Rodrigo-Carranza et al., 2021, 2023a; Senefeld et al., 2021). RE is commonly defined as the steady-state oxygen uptake or metabolic energy required at a given submaximal speed (Barnes & Kilding, 2015). Changes in RE translate directly into changes in running performance (Hoogkamer et al., 2016; Kipp, Kram & Hoogkamer, 2019). However, in track events athletes typically run at an intensity far above their aerobic steady-state capacity (except for during the 10,000m) (Brandon, 1995; Alvero-Cruz et al., 2020), and therefore RE measures cannot be assessed at race intensity. This is particularly relevant for spikes, since benefits from footwear are likely to be speed-specific (Mcleod et al., 2020).

To overcome this challenge, we recently introduced and validated a novel approach to assess the benefits of AFT spikes, consisting of a series of 200m trials at self-perceived middle-distance race pace on the track (Bertschy et al., 2024). Between this interval-based protocol for middle-distance track events (800/1500m), time trials (3,000/5,000m) and RE measurements for long-distance track events (10,000m), tools are now available to assess the benefits of AFT spikes, specific to each event.

Sprint spikes with increased longitudinal bending stiffness have been around since before AFT shoes and have been shown to improve top speed (Stefanyshyn & Fusco, 2004; Willwacher et al., 2016), although with mixed results (Smith et al., 2016; Nagahara, Kanehisa & Fukunaga, 2017). Willwacher et al. (2016) showed that how increased longitudinal bending stiffness affects sprint outcomes depends on an individual’s plantar-flexor strength. Further, the optimal amount of bending stiffness might be speed dependent (Roy & Stefanyshyn, 2006; Mcleod et al., 2020; Day & Hahn, 2020). Still, AFT road shoes with a carbon fiber plate have been shown to be effective at speeds as slow as 3.89 m/s (Hoogkamer et al., 2018), and almost all competitive AFT marathon shoe models have a plate or other stiffening elements (Joubert & Jones, 2022). This suggests that AFT spikes (used at speeds beyond 5.5 m/s) will be most effective *with* stiffening elements. However, the exact function of carbon fiber plates within AFT marathon shoes is still under debate (Nigg, Cigoja & Nigg, 2021; Kram, 2022; Rodrigo Carranza, 2023), with increasing evidence that the improved foam properties are most important (Hoogkamer, Kipp & Kram, 2019; Healey & Hoogkamer, 2022; Kram, 2022). Indeed, race results suggest that for long-distance events AFT spikes with modern foam *without* a plate might be as effective as AFT spikes that combine modern foam *with* a plate. To specifically address this, we compared two types of AFT spikes against traditional spikes, at speeds specific to various middle-distance and long-distance track events.

Existing research on AFT spikes is scarce. Oehlert et al. (2023) showed that two different commercially available AFT spike models (with and without elements to increase the longitudinal bending stiffness) improved RE by ∼2% compared to a traditional spike model, during treadmill running at 4.44 m/s. For track running at ∼6.5 m/s, Bertschy et al. (2024) found that participants ran 2-3% faster in different prototype and commercially available AFT spike models than in traditional spikes. The increased speeds in the AFT spikes were accompanied by longer steps, with no significant changes in step frequency. On the other hand, in a case study Russo et al. (2022) evaluated the biomechanics of running with AFT spikes and traditional spikes on the track at ∼8.0 m/s and found no changes in step length or step frequency, but found longer contact times in the AFT spikes.

The aim of the current study was to examine the influence of different AFT spike components on middle-and long-distance performance measures and spatiotemporal variables in trained runners. We hypothesized that at middle-distance pace AFT spikes combining a modern foam with a curved carbon fiber plate would have the largest improvement in running speed, similarly in females and males. We hypothesized that at long-distance pace AFT spikes with modern foam without a plate would have similar RE as AFT spikes that combine a modern foam with a curved carbon fiber plate, but better RE compared to traditional spikes. Small spatiotemporal modifications such as a reduction in step frequency and an increase in contact time were expected for running in AFT spikes compared to in traditional spikes. Additionally, we evaluated perceived comfort in and perceived performance enhancement from AFT spikes (Hébert-Losier et al., 2024).

## MATERIALS AND METHODS

### Participants

Fourteen trained female runners (mean ± SD: 21.3 ± 4.2 years; 50.9 ± 3.5 kg and 164 ± 4 cm) and sixteen trained male runners (27.1 ± 4.9 years; 63.3 ± 4.3 kg and 176 ± 5 cm) participated in this study. The inclusion criteria were the following: 1) running for at least five days per week during the previous six months without any injury, 2) ability to run 10 km in less than 40 min (females) or 34 min (males), 3) fitting a women’s EU38 or a men’s EU42–44 shoe size and 4) regularly running in spikes and AFT shoes. Prior to the study, all participants were informed about the testing protocols and possible risks involved, and were invited to provide written informed consent. We recruited participants from Castilla La-Mancha and Madrid during the outdoor track season by contacting their coaches or contacting them directly. The study was performed in accordance with the principles of the Declaration of Helsinki (October 2008, Seoul), and the experimental protocols were approved by the local ethics committee (CEIC924).

### Spike characteristics

A traditional spike model with an ethylene vinyl acetate (EVA) midsole, without a carbon fiber plate served as our control condition (“Control”, Puma evoSPEED Mid-Distance). This was compared against an AFT spike model with modern foam (“PEBA”, Puma evoSPEED Long Distance Nitro Elite+) and against an AFT spike model with modern foam and a curved carbon fiber plate (“PEBA+Plate”, Puma evoSPEED Distance Nitro Elite+) (Figure 1). All spikes had standardized 6 mm spike pins for traction. The average mass was 158g for the control and 151g and 149g for the PEBA and PEBA+Plate conditions, respectively, for EU42 (Table 1). Participants were able to see the spikes, but they could not manipulate the spikes and were not aware of any differences in spike properties.

**Figure 1.**
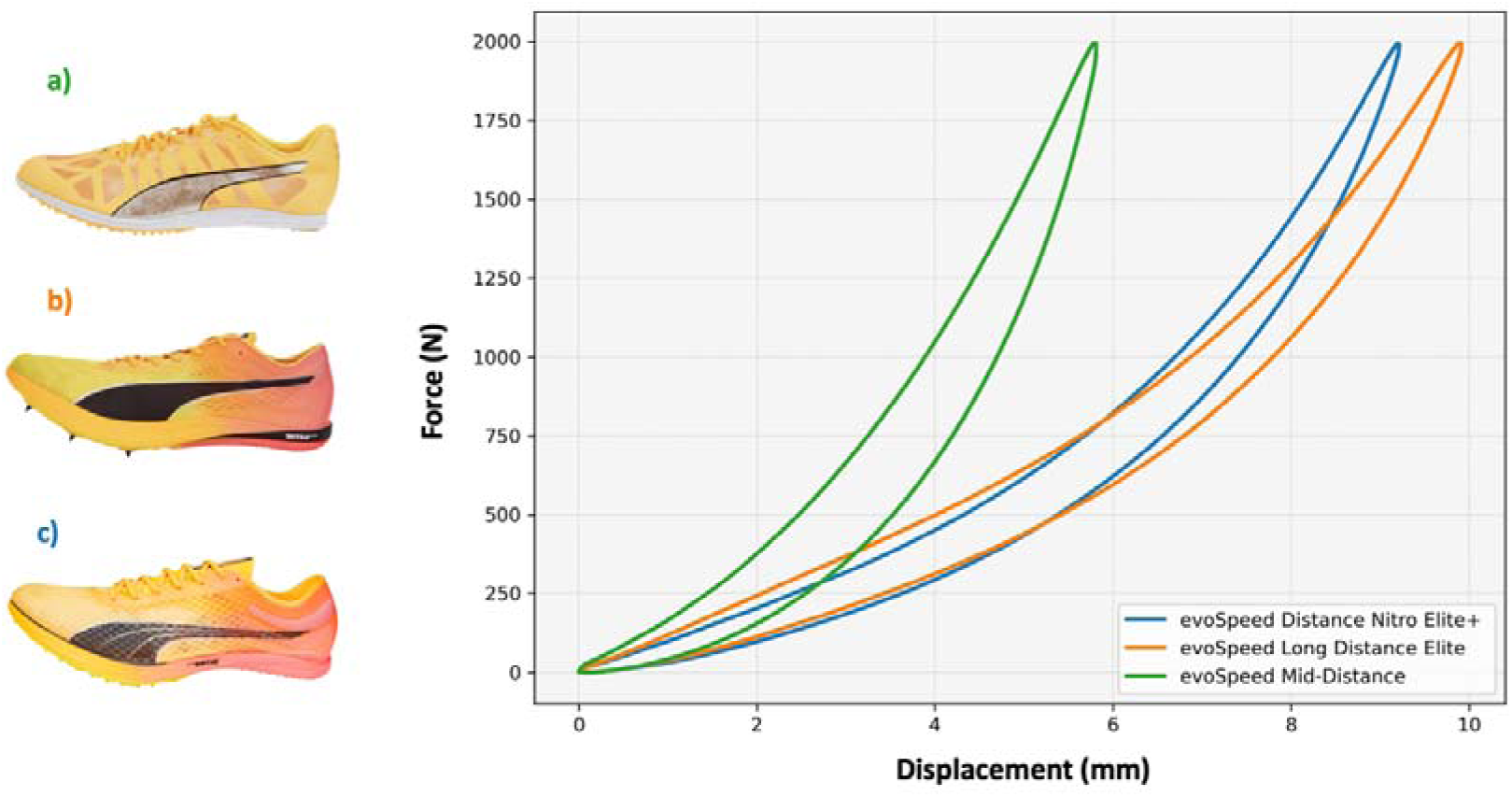
Spike conditions. a) Control: Traditional spike (evoSPEED Mid-Distance), b) PEBA: modern midsole, no plate (evoSPEED Long Distance NITRO Elite+), c) PEBA + Plate: modern midsole + carbon fiber plate (evoSPEED Distance NITRO Elite).

**Table 1.** Characteristics of experimental spike conditions.

| Characteristics | Control | PEBA | PEBA+Plate |
| --- | --- | --- | --- |
| Stiff plate | No | No | Full length |
| Midsole foam | EVA | PEBA | PEBA |
| Mass (g) | 158 | 151 | 149 |
| Midsole thickness (mm) | 12 | 20 | 18 |
| Maximal deformation (mm) | 5.8 | 9.9 | 9.2 |
| Overall stiffness (N/mm) | 344.2 | 201.8 | 217.2 |
| Energy returned (%) | 67.2 | 77.1 | 78.2 |
| Spike characteristics for EU42 |  |  |  |

The force-displacement relationship of the spikes was determined by a compression test on the complete spike (including shoe upper) where a custom-made structure simulating a size EU42 foot was attached to a material testing machine that measures force and displacement (Zwick/Roell, Ulm-Einsingen, Germany). A force of 2000 N was applied at 6 Hz (0.167 s) to simulate the running velocities evaluated in this study. Cycle 99 of 105 was analyzed after ensuring that displacement and force amplitude were consistent across previous cycles. Overall stiffness was calculated as the maximum force (2000 N) over the maximum deformation. The measurement was repeated on 3 different days showing a very good inter-day variability.

### Design and methodology

We evaluated the effects of AFT spikes on middle- and long-distance running performance measures using a randomized mirrored experimental design. All participants refrained from intense exercise in the 48h preceding the test sessions. Test sessions were performed on an outdoor athletics track (400 m) with controlled environmental conditions (550 m altitude, 22-25°C, 26-29% relative humidity). Participants were asked to avoid caffeine and alcohol intake within 24 hours of their testing visits and refrained from eating and drinking anything but water for 4h before testing.

The female runners completed one single visit. Following the novel interval-based approach to evaluate middle-distance spikes of Bertschy et al.,(2024) participants performed six experimental trials of 200m, controlling effort at self-perceived 800m race pace. After a standardized warm-up of 10 min of jogging and two runs of 200m at a pace close to their 800m race pace (to ensure that the pace was constant), the participants completed six 200m trials, two in each spike condition, in a mirrored order (a-b-c-c-b-a). Participants were unaware of the times they were running for each trial as they were not using a watch and researchers did not disclose the times they recorded until after the data collection was complete. Rest between trials was 8 minutes, to allow for changing spikes, enough rest and to go to the start of the next trial. Participants put on the spikes and ran 100m in the direction of the starting line to test each spike, prior to the trial. They then walked the final 100m to the start line to avoid fatigue. They then ran the 200m trial from a standing start (similar to a middle-distance race start) which was timed by two researchers. For the timing of each trial, researchers started their timers when the participant started running, indicated by the participant quickly lowering one hand from a raised position, and stopped their stopwatches when the participant crossed the finish line. Subsequently to the 200m trials, the female runners performed three trials of 800m at 4.44 m/s (16 km/h), one in each spike condition, with 8 min rest between trials, with the objective of measuring running biomechanics at a constant speed. We chose 4.44 m/s according to the level of the participants, targeting a metabolic steady state speed, with the purpose to evaluate a long-distance running pace. During the 800m trials the speed was controlled by sound signals every 22.5 seconds. The participants aimed to cover 100m between subsequent sound signals, which coincided with running 4.44 m/s.

The male runners performed four different visits separated by five days. During the first three visits the participants completed six experimental trials of 200m controlling effort at self-perceived 800m race pace with the same protocol as the female runners (Bertschy et al., 2024). After this, the male runners performed a 3,000m time trial wearing one of the three spike conditions. The participants received verbal information about the number of laps completed and to go, but no other feedback or encouragement was provided during the test. Participants were kept blind to time during each of the 3,000m time trial test. The sequence of the shoes was randomized for each participant using a random number generator. We quantified pacing by expressing each lap as a percentage of the average 3,000m time trial speed.

The male runners performed an additional, fourth, visit consisting of a series of 3-lap RE trials at 5.00 m/s (18 km/h; i.e., 4-minute trials) with each condition twice, in a mirrored order (total 2 × [3 × 4 min]). During the 4-minute trials, the speed was controlled by sound signals every 20 seconds. The participants aimed to cover 100m between consecutive sound signals, which coincided with running 5 m/s. We chose 5 m/s so the participants should be able to run below the second ventilatory threshold to ensure steady-state V[O_2_ measurements. This was confirmed by respiratory exchange ratio measurements (values below 1.0). During the RE trials, we measured respiratory variables using a PNOE gas analyzer (ENDO Medical, Palo Alto, CA, USA)(Tsekouras et al., 2019) which was calibrated prior to each session (CO_2_ 4.10%; O_2_ 15.92%). We used V[O_2_ and V[CO_2_ values, collected during the last minute of each 4-minute trial, to calculate RE in W/kg (Peronnet & Massicotte, 1991; Saunders et al., 2004; Kipp, Byrnes & Kram, 2018). Each participant performed all of their running trials on the same athletic track, but not all participants ran on the same track (five different tracks in total). Recruiting a sample of 14 female runners, meeting our inclusion criteria proved to be challenging and resulted a large geographical spread within this group. Travel and time limitations (outdoor season was over for most of the participants) did not allow us to have the female runners execute all four visits that the male runners performed. So, we adapted the protocol to a single visit for the female runners.

We measured the main spatiotemporal parameters (contact time and step frequency) for every step during the self-paced 200m trials and the controlled speed 800m trials for the female runners, and during the self-paced 200m trials, the 3,000m time trial and the controlled speed RE trials for the male runners, using a Stryd Power Meter device (V2, Stryd, Boulder, CO, USA), sampling at 1000 Hz (Rodríguez-Barbero et al., 2024). The Stryd device was paired with a Polar® watch (Vantage M GPS system; Polar, Kempele, Finland) and the information was analyzed in the web-based Stryd Power Center. In the 200m trials, the spatiotemporal parameters were extracted for 50-150m (at least thirty steps). Twenty-five steps provide enough reliability for running mechanics measurements to distinguish running technique between people (Oliveira & Pirscoveanu, 2021). For the 800m trials for female runners and the 4-minute RE trials for male runners we evaluated the central minute of each trial. During the 3,000m time trial, the spatiotemporal variables were measured during the whole test and were averaged for subsequent analyses.

At the end of each 200m trial at self-perceived 800m race pace, and each long-distance running pace trial (800m at 4.44 m/s for female runners, 4 min at 5 m/s for male runners) in each spike condition, participants provided their perception of spike comfort and performance enhancement on a subjective scale (Lindorfer, Kröll & Schwameder, 2019) from 0 to 100, and then rested before starting the next trial. The questions were, “how comfortable were the shoes?” and “how much do you think the shoes help you during running?” The corresponding anchor points for these scales were 0 = “Least comfortable spike I have ever worn” to 100 = “Most comfortable spike I have ever worn”, and 0 = “Least helpful spike I have ever worn” to 100 = “Most helpful spike I have ever worn”, respectively.

### Statistical analysis

Statistical analyses were carried out using Jamovi Statistics for Macintosh, Version 2.4.1 (Jamovi project, Boston, MA, USA). We assessed all outcomes of the 200m trials at self-perceived 800m race pace and spatiotemporal variables at long distance pace with a linear-mixed effect model (LMEM) (Wilkinson, Mazzo & Feeney, 2023). The LMEM had spike condition and sex as fixed effects, participant as a random intercept and speed and spatiotemporal variables as the output variables. Additionally, we used LMEM for all outcome variables of the 3,000m time trials (speed, stride frequency and contact time) and RE trials with spike condition as fixed effects and participant as a random intercept. For the 3,000m time trials, we evaluated the pacing (% of average speed, for each lap) using a LMEM with spike condition and lap as fixed effects, participant as a random intercept and speed as the output variable. Pairwise comparisons with Bonferroni adjustment were used when any significant main or interaction effect was identified. The sphericity assumption was checked with Mauchly’s test followed by the Greenhouse-Geisser adjustment where required. Results are expressed as mean ± standard deviation. We used Pearson correlation coefficients to investigate the relationship between the improvements in the 200m trials at self-perceived 800m race pace and the improvements in the 3,000m time trials for each of the AFT spike models vs. Control. The significance level was set at p < 0.05 for all analysis.

## RESULTS

### Self-perceived 800m race pace

There was a significant effect for spike condition (p<0.001) and spike × sex interaction (p=0.03) on running speed for the self-perceived 800 m race pace trials. Females ran significantly faster in PEBA (5.58±0.59 m/s; p=0.003; 2.1%) and PEBA+Plate (5.57±0.56 m/s; p=0.001; 2.0%) compared to Control (5.46±0.58 m/s). Males ran significantly faster in PEBA (6.56±0.31 m/s; p<0.001; 1.4%) and PEBA+Plate (6.63±0.36 m/s; p<0.001; 2.4%) compared to Control (6.47±0.33 m/s), but males also ran significantly faster in PEBA+Plate than in PEBA (p=0.002; 1.1%) (Figure 2).

**Figure 2.**
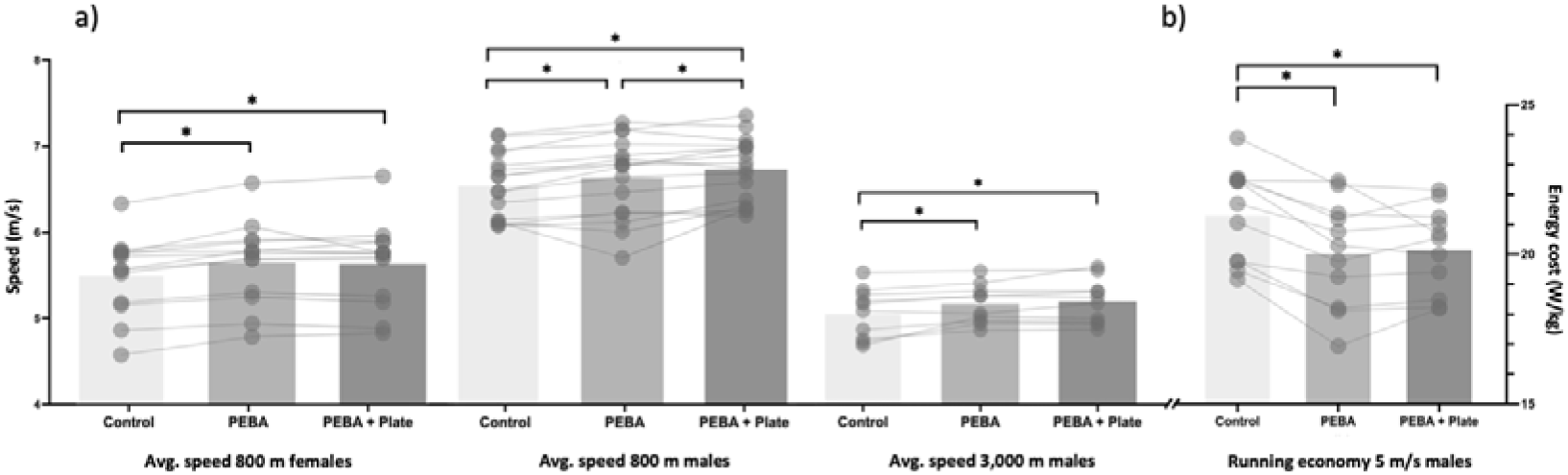
a) Average 800 m and 3,000 m running speed for female and male runners; b) running economy (W/kg) at 5 m/s for male runners. Bar graphs represent mean values and circles represent runners. \**p* ≤ 0.05 during *post hoc* comparisons when main effect of spike condition was significant.

In the 200m trials at self-perceived 800m race pace, there was a significant effect for spike condition on step frequency (p<0.001) and contact time (p=0.002; Table 2). For females, average step frequency was significantly lower in PEBA (195.2±13.9 step/min; p<0.001; 1.7%) and PEBA+Plate (194.1±15.8 step/min; p<0.001; 1.6%) than in Control (198.6±15.4 step/min). For males, average step frequency was significantly lower in PEBA+Plate (197.4±11.6 step/min; p=0.001; 2.4%) than in Control (202.1±11.3 step/min) but without significant differences vs. PEBA (200.3±10.6 step/min). Contact time was significantly longer for males in PEBA+Plate (0.189±0.029 s) than in Control (0.176±0.024 s; p=0.010; 7.4%), but similar between conditions for the female runners.

**Table 2.**
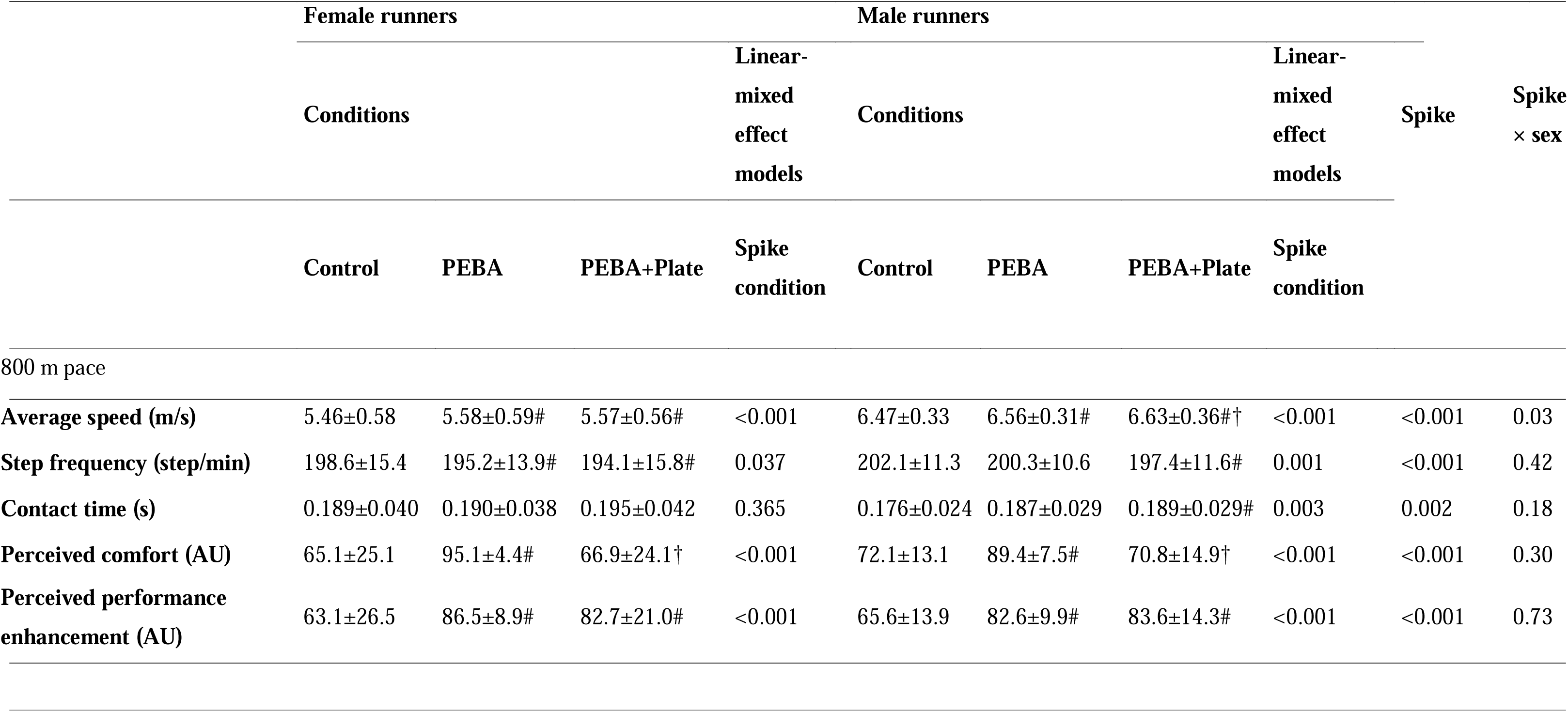
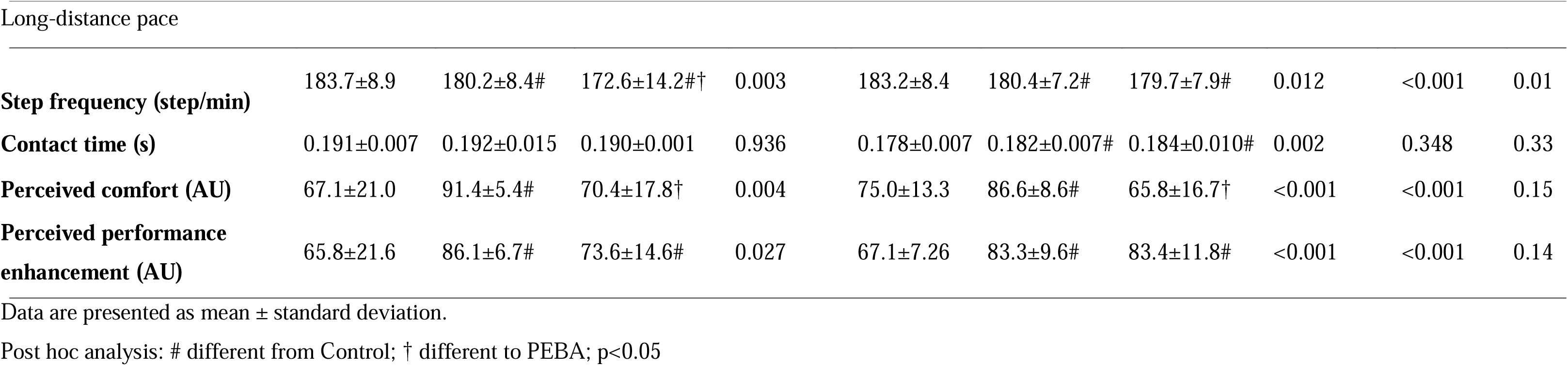
Results of spatiotemporal variables during the trials at 800 m race pace and at long-distance pace for each spike condition for female nd male runners.

### 3,000m time trials

Four male runners did not complete at least one of the 3,000m time trials due to personal reasons and were eliminated from the analysis. There was a significant main effect of spike condition on speed during the 3,000m time trial (p=0.013, n = 12; Table 3). Speed improved in PEBA (5.79±0.25 m/s; p=0.034; 1.0%) and PEBA+Plate (5.87±0.37 m/s; p=0.03; 2.4%) compared to Control (5.73±0.29 m/s). The main effect for spike condition on pacing was not significant (p=0.458).

**Table 3.** Results for performance and submaximal running variables for each spike condition for males.

|  | Conditions |  |  | Linear-mixed effect models |
| --- | --- | --- | --- | --- |
|  | Control | PEBA | PEBA+Plate | Spike condition |
| **Running Economy at 5 m/s (W/kg)** | 21.36±1.51 | 20.25±1.23# | 20.47±1.30# | <0.001 |
| **Speed during 3,000 m time trial (m/s)** | 5.73±0.29 | 5.79±0.25# | 5.87±0.37# | 0.011 |
| **Step frequency during 3,000 m time trial (step/min)** | 188.7±7.1 | 187.2±7.5 | 186.1±7.8# | 0.041 |
| **Contact time during 3,000 m time trial (s)** | 0.180±0.007 | 0.181±0.013 | 0.180±0.005 | 0.866 |

Data are presented as mean ± standard deviation.
Post hoc analysis: # different from Control, p<0.05

There was a significant main effect of spike condition on step frequency during the 3,000m time trials (p=0.041) with significantly lower step frequency in PEBA+Plate (186.1±7.8 step/min; p=0.03; 1.4%) compared to Control (188.7±7.1 step/min), but PEBA was not significantly different (187.2±7.5 step/min). The main effect for spike condition on contact time (p=0.866) was not significant.

Regression analysis showed that the speed improvements from the control condition for the 200m at self-perceived 800m race pace and for the 3,000m time trial were strongly correlated for both PEBA (r=0.891; p<0.001) and PEBA+Plate (r=0.832; p<0.001). In the 200m trials at self-perceived 800 m race pace 63% of the participants (females and males) ran fastest in PEBA+Plate and 37% of the participants ran fastest in PEBA. In the 3,000 m time trials, 59% of the participants (males only) ran fastest in PEBA+Plate, 33% of the participants ran fastest in PEBA, and one participant ran fastest in Control (8%).

### Running Economy trials

Four male runners did not complete the RE trials due to respiratory exchange ratios greater than 1.0 (n = 2) or personal reasons (n = 2) and were eliminated from the analysis. During the RE assessment session, respiratory exchange ratio remained below 1.0 across all trials for all participants included in the study (n=12) (0.92 ± 0.11 in the final minute). Moreover, there was no indication of any substantial V[O_2_ and V[CO_2_ slow component, as oxygen uptake and carbon dioxide production between minute 3 to 4 differed by less than 0.5% (p=0.86; p=0.79).

There was a significant effect of spike condition on RE (p<0.001). RE was significantly better in PEBA (20.25±1.23 W/kg; p<0.001; 5.1%) and PEBA+Plate (20.47±1.30 W/kg; p<0.001; 4.0%) than in Control (21.36±1.51 W/kg) (Figure 2).

### Controlled speed spatiotemporal parameters

Regarding spatiotemporal parameters during the long-distance pace trials (4.44 m/s for females and 5 m/s for males) there was a significant effect of spike condition on step frequency (p<0.001) and spike × sex interaction (p=0.01; Table 2). In females, step frequency was significantly lower in PEBA+Plate (172.6±14.2 step/min) than in PEBA (180.2±8.4 step/min; p<0.005; 4.0%) and Control (183.7±8.9 step/min; p<0.005; 5.8%), moreover, PEBA+Plate was significantly lower than PEBA (p<0.005). In males, step frequency at long-distance pace was significantly lower in PEBA (180.4±7.2, p=0.004; 1.6%) and PEBA+Plate (179.7±7.9 step/min, p<0.001; 1.9%) than in Control (183.2±8.4 step/min). The main effect of spike condition on contact time was not significant (p=0.348).

### Subjective variables

Regarding subjective variables, there was a significant effect of spike condition on perceived comfort (p<0.001) and perceived performance enhancement (p<0.001) during the self-perceived 800m race pace trials, and perceived comfort (p<0.001) and perceived performance enhancement (p<0.001) during the long-distance pace trials. There was no significant spike × sex interaction for any of the subjective variables. Both females and males perceived PEBA to be more comfortable than Control and PEBA+Plate at both 800m race pace and long-distance pace (Table 2). Between subject variability was larger in PEBA+Plate than in Control and PEBA across sex and speed conditions. Both females and males perceived both PEBA and PEBA+Plate to be more performance enhancing than Control at both 800m pace and long-distance pace (Table 2).

## DISCUSSION

The aim of this study was to examine the influence of different AFT spike components on middle- and long-distance performance measures and spatiotemporal variables in trained runners. Females and males both ran faster at self-perceived 800m race pace in PEBA (2.1% and 1.4%, respectively) and PEBA+Plate (2.0% and 2.4% respectively) compared to Control. Moreover, males ran significantly faster in PEBA+Plate than in PEBA (1.1%). In the 3,000m time trial, males ran 1.0% faster in PEBA and 2.4% faster in PEBA+Plate. Speed improvements in the 3,000m time trial had a large correlation with the improvements at self-perceived 800m race pace. Further, RE improved by 5.1% and 4.0% in PEBA and PEBA+Plate compared to Control. The improvements in average speed during the self-perceived 800m race pace trials and during the 3,000m time trial and in RE were accompanied by spatiotemporal modifications, mainly a reduction in step frequency. Participants perceived PEBA to be more comfortable than Control and PEBA+Plate, and perceived both AFT conditions to be more performance enhancing than the control spikes.

### Performance variables

Our results show that females and males improved similarly with AFT spikes compared to control, although females improved slightly more than males when using PEBA (2.1 vs. 1.4%). Moreover, males ran faster in PEBA+Plate than in PEBA while there was no difference for females. Barnes and Kilding (2019) found similar RE improvements in females and males running in AFT road shoes (with plate) compared to traditional spikes and to traditional marathon road shoes for running speeds slower than those observed in our study (3.9 and 5.0 m/s). On the other hand, Mason et al.,(2023) demonstrated that female sprint events have undergone more substantial and more widespread recent improvements than male events, similar to observations on road running results (Bermon et al., 2021). Our results show that the performance improvements at 800 m speed are similar in females and males, but that females might benefit more from PEBA spikes without plate than males, and that males benefit more from PEBA+Plate spikes than from PEBA spikes without plate. The higher running speed of males (∼6.5 m/s) compared to females (∼5.6 m/s) and anthropometric differences such as females being shorter (∼165 vs. ∼175 cm) and lighter (∼50 vs. ∼65 kg) could have affected this, since optimal longitudinal bending stiffness appears to depend on body mass (Roy & Stefanyshyn, 2006) and running speed (Day & Hahn, 2020; Rodrigo-Carranza et al., 2023b) and may differ between athletes with different strength abilities (Willwacher et al., 2016).

Regarding the 3,000m time trials, our results provide novel evidence that AFT spikes elicit better running performance than traditional spikes, while running at intensities higher than those measured in the RE tests. The improvements of 1.0 and 2.4% using PEBA and PEBA+Plate spikes, respectively, are in the same order as those reported in retrospective studies that analyzed the progression of race times using public data in relation to the launch of the first AFT spikes ranging from 0.5 to 2% for sprint events up to 400 m (Mason et al., 2023) and middle-distance events (Willwacher et al., 2024). The observed between-spike differences in the 3,000m time trials were very similar to those observed in the 200m trials at self-perceived 800m race pace. At 800m race pace PEBA was 1.4% better than Control as compared to 1.0% better in the 3,000m time trials (r=0.891, p<0.001), while PEBA+Plate was 2.4% better than Control in both events (r=0.832, p<0.001). These observations provide additional external validity for interval-based approach using 200m trials at self-perceived race pace recently introduced by Bertschy et al.(2024). That the difference between PEBA and PEBA+Plate was only significant at self-perceived 800m race pace could be due to the difference in running speed, or because participants performed more separate trials at 800m pace (providing more statistical power). Combined, our results suggest that both AFT spike technologies improve middle-distance track performance, and that spikes with increased longitudinal bending stiffness perform better than spikes only using PEBA.

To assess the effects of AFT spikes on long-distance running performance, we measured RE at a fixed speed of 5 m/s. In-line with the faster 3,000m time trials in the AFT spikes, RE improved in the AFT spikes. RE improved 5.0 and 4.1% beyond Control in PEBA and PEBA+Plate, respectively. These RE improvements are as substantial as those repeatedly reported for AFT road shoes vs. traditional racing flats (Hoogkamer et al., 2018; Barnes & Kilding, 2019; Whiting, Hoogkamer & Kram, 2022). The relation between improvements in submaximal RE and time trials performance has been shown before (Hoogkamer et al., 2016) and theoretical modeling of the relationship between RE and running performance (Kipp, Kram & Hoogkamer, 2019) suggests that at speeds beyond 3 m/s time trial improvements will be smaller than RE improvements. Statistically the 3,000m time trial results and the RE results are in line, with both AFT spike models performing better than the control spikes, without significant differences between the AFT spike models. While not significant, it is noteworthy that group average values showed 1.4% faster time trials, but 1.1% worse running economy in PEBA+Plate than PEBA. Altogether this suggests that AFT spikes outperform traditional spikes across all middle-and long-distance events, and that AFT spikes with plate are better than AFT spikes without plate for the 800m, but that between-spike differences at longer track distances are less clear, differ between individuals, and are speed dependent: the average speeds across the different tests were 6.5 m/s (800m race pace), 5.7 m/s (3,000m time trials) and 5.0 m/s (RE trials) in the control spikes for the males.

### Spatiotemporal modifications

Spatiotemporal changes for running with AFT have been investigated by many researchers (Hoogkamer et al., 2018; Hoogkamer, Kipp & Kram, 2019; Barnes & Kilding, 2019; Ortega et al., 2021) with changes in step length and contact time being observed most commonly (Hoogkamer et al., 2018; Hunter et al., 2019). In the current study, we found a reduction in step frequency at all speeds evaluated (4.5-6.5 m/s) for both AFT spike conditions, except for male runners in PEBA at self-perceived 800 m race pace. The reductions in step frequency with AFT spikes are particularly noteworthy in relation to the increased running speeds. Although step length was not available from the measuring device used in the study, average step length can be calculated from average speed and average step frequency. Using this, average step length increased 6 cm in PEBA and PEBA+Plate vs. Control at self-perceived 800m race pace in female runners (∼1.71 m for both AFT spikes vs. ∼1.65 m; 3.6%). For male runners, average step length increased 3 cm in PEBA (∼1.95 m; 1.0%) and 9 cm in PEBA+Plate (∼2.01 m; 4.7%) vs. Control (∼1.92 m). Participants experienced a similar reduction in step frequency when running with AFT spikes in the 3,000m time trial. Over the full distance, participants took ∼34 steps less in PEBA vs. Control (∼1651 vs. ∼1617 steps) and 66 steps less in PEBA+Plate vs. Control (∼1651 vs. ∼1585 steps), which coincides with a ∼4 cm longer step length in PEBA and a ∼7 cm longer step length in PEBA+Plate, than in Control. Similarly, in the constant speed trials (4.44 m/s for females and 5.00 m/s for males), females increased step length by ∼3 cm in PEBA (∼1.48 m; 2.1%) and ∼9 cm in PEBA+Plate (∼1.54 m; 6.2%) vs. Control (∼1.45 m) and males increased step length by ∼2 cm in PEBA (∼1.66 m; 1.2%) and ∼7 cm (∼1.71 m; 4.1%) vs. Control (∼1.64 m).

Previous studies have reported similar step frequency decreases (1.3-1.9%) (Hoogkamer, Kipp & Kram, 2019; Hata et al., 2022) and proportional step length increases (1.2-1.4%)(Hunter et al., 2019; Barnes & Kilding, 2019; Hata et al., 2022) in AFT road shoes compared to traditional running shoes during treadmill running at constant speeds (Hunter et al., 2019; Barnes & Kilding, 2019; Hata et al., 2022; Joubert, Dominy & Burns, 2023), during track running in AFT road shoes (1.3-2.7%) (Hébert-Losier et al., 2024) and recently in AFT spikes (1.9-2.3%) (Bertschy et al., 2024). However, the increases in step length we observed here in PEBA+Plate (4.1-6.2%) are substantially larger than those reported before.

In our study, females and male showed similar step frequency changes in AFT spikes at self-perceived 800m race pace. However, males ran with significantly increased contact time in PEBA+Plate compared to Control, while females did not significantly increase contact time (displaying more interindividual variability and without spike × sex interaction). Russo et al., (2022) also observed longer contact times in AFT spikes with a plate vs. traditional spikes in a case study of a female athlete running at ∼8.0 m/s. The increased contact time in PEBA+Plate compared to Control could be related to a slower, longer propulsion phase (Willwacher et al., 2014) related to the increased longitudinal bending stiffness of PEBA+Plate (Ortega et al., 2021; Rodrigo-Carranza et al., 2022).

### Subjective variables

Participants perceived the PEBA and PEBA+Plate spikes to be more performance enhancing than Control, similarly in females and males running at self-perceived 800m race pace and at long-distance race pace. However, PEBA scored higher in comfort than Control and PEBA+Plate. This is similar to observations by Hébert-Losier et al.(2024) comparing an AFT road shoe to a minimalist road shoe (which in several aspects might not be too different from a traditional spike). This increased comfort in PEBA might have additional benefits in long-distance track races (i.e., 10,000m) for which comfort might have increased relevance.

## LIMITATIONS

The present study has several limitations. First, the 3,000m time trial performance test and the RE test were performed only by the male runners. Therefore, although the females and males had similar performance and spatiotemporal results for the 200m trials at self-perceived 800m race pace, the improvements in the 3,000m time trial and in RE might not generalize to female runners. In addition, in the 200m trials at self-perceived 800m race pace, runners simply could have run faster because they perceived the AFT spikes to be more performance enhancing. However, the high correlation for between-spike differences between the 200m trials and the 3,000m time trials, combined with the results from the added mass experiment by Bertschy et al. (2024) suggest this is not the case. Although the effects of variability in the performance tests were reduced by including multiple trials in the same spikes in the analyses (Bertschy et al., 2024) and by using two RE measurements in each condition and in counterbalanced order within a single session (Barrons et al., 2024), the 3,000m time trials were only performed once in each condition and these results might be less robust.

## CONCLUSIONS

This is the first study that compares the effects of different components of AFT spikes on middle-and long-distance running performance measures. Confirming our hypothesis, both AFT spike technologies improved middle-and long-distance running performance measures. At self-perceived 800m race pace females ran 2.1 and 2.0% faster in PEBA and PEBA+Plate spikes as compared to traditional spikes, and males ran 1.4% faster in PEBA spikes and 2.4% faster in PEBA+Plate spikes. In the 3,000m time trials, males ran 1.0% faster in PEBA spikes, while they ran 2.4% faster in PEBA+Plate spikes, each compared to traditional spikes. Furthermore, running economy was 5.1% and 4.0% better in PEBA and PEBA+Plate spikes, respectively, compared to traditional spikes. The improvements in running performance and running economy were accompanied by small modifications in running kinematics, mainly a reduction in step frequency, and by an increase in perceived performance enhancement in both AFT spikes compared to traditional spikes. Overall, the results of this study indicate that footwear companies should design AFT spikes that combine PEBA foam with a carbon fiber plate to improve performance in middle-distance events for males. However, for middle-distance events for females and for longer events for males, both spike conditions seem equivalent based on short-term performance measures. The increased perceived comfort in the PEBA spikes suggests this could be the best option for long-distance track events, particularly the 10,000m.

## Compliance with Ethical Standards

### Acknowledgements

We thank the participants for participating, PUMA SE for providing the spikes for the study, Montgomery Bertschy for his help with the statistical analysis and Rory Tatterton for his help with the mechanical testing.

### Ethical approval

The study was performed in accordance with the ethical standards of the Declaration of Helsinki. Ethics approval was obtained from the University of Castilla La-Mancha Institutional Review Board (CEIC924).

### Authors’ contributions

Conceptualization: Victor Rodrigo-Carranza and Wouter Hoogkamer; Experimental design: Victor Rodrigo-Carranza, Wouter Hoogkamer; Data collection: Victor Rodrigo-Carranza, Violeta Muñoz-de la Cruz; Data analysis: Victor Rodrigo-Carranza and Wouter Hoogkamer; Writing - original draft preparation: Victor Rodrigo-Carranza, Violeta Muñoz de la Cruz, Wouter Hoogkamer; Writing - review and editing: Victor Rodrigo-Carranza, Wouter Hoogkamer

